# Effects of a novel three-dimensional grid intrauterine device on the uterus, steroid receptor and PAX2 of rhesus macaques

**DOI:** 10.1101/858886

**Authors:** Mei-Hua Zhang, Li-Ping Zhai, Ling Yu, Xia Song, Jian-Chun Yu, Yi Qiu

**Author notes:** These authors contributed equally to this work (Mei-Hua Zhang, Li-Ping Zhai). Corresponding author: Jian-Chun Yu, Yi Qiu.

## Abstract

Intrauterine devices (IUDs) is the most effective methods of the reversible and long-acting contraception. 1) To develop a novel three-dimensional grid intrauterine device (3-DGIUD) with nickel-titanium (Ni-Ti) and silicone rubber. 2) To observe the effect of the 3-DGIUD on contraceptive efficacy and the change of uterus, endometrial sex steroid receptor, PAX2 in rhesus macaques (Macaca mulatta). The materials of the 3-DGIUD were the nitinol wire and the silicone rubber. The frame of the 3-DGIUD was three-dimensional and grid-like. Twenty adult female rhesus macaques were divided into the 3-DGIUD group (placing the 3-DGIUD, n=9), the sham operation group (no placing the 3-DGIUD, n=9) and the control group (n=2). On the 10^th^-day after surgery, the 3-DGIUD group and the sham operation group macaques were caged together with male macaques (female: male = 1:1). The uterus, 3-DGIUD and pregnancy of 18 female rhesus monkeys were examined by abdominal ultrasound every month. The endometrium pathological examination was carried out and the expression of PAX2 and hormone receptor (ER, PR) was detected by immunohistochemical staining. After 3-DGIUD was placed in case group for 3 and 12 months, only 1 of female macaque was pregnant in 9. The contraceptive effective rate was 88.9% (8/9). The 3-DGIUD in the uterus of macaques was observed by ultrasound. In the sham operation group, 9 macaques were pregnant (9/9). There was significant difference in uterine size of the 3-DGIUD group between pre-placement and after surgery for 3 and 12 months (*P*<0.05). The endometrial epithelium was intact, just a small number of glands vacuoles and a few neutrophils infiltration around the 3-DGIUD. The expression of endometrial ER, PR and PAX2 in 3-DGIUD group on 12 months after surgery was similar to those in control macaque. The 3-DGIUD has a good contraceptive effect on female macaques, and has no significant affection on the expression of endometrial steroid receptor and PAX2 in rhesus monkeys.

## Introduction

Appliance of contraceptives is the cornerstone of prevention for unintended pregnancy. The rate of unintended pregnancy in the United States is 51% and 45% in 2011 and in 2008, respectively [1]. In China, more than 20% of teenager and unmarried women are pregnant every year [2], and the rate of undergoing repeat abortions and non-use of contraception in these adolescents is 39% and 68%, respectively [3]. Long-acting reversible contraceptive (LARC) methods could help reduce the high rate of unintended pregnancy. Among all available contraceptive methods, the failure rate of IUD is less than 1%, which is almost ideal and the most popular contraceptive method in the world [4, 5]. IUDs are the most common forms for millions of women, particularly in mediate postpartum women [6–8]. Current data shows that more than 10% of childbearing aged women are using IUDs worldwide [4]. In China, IUDs are utilized by 48%-51% women of childbearing aged in 2010-2015 [5, 8]. The rate of IUD use in American Muslim women are 21.2% [9]. In breast cancer women, the rate of IUDs use is 72.1% [10].

Although female contraceptive methods such as sterilization, IUDs and hormonal contraception are very effective in preventing unintended pregnancy, some women can’t use them due to health condition or side effects. IUDs may cause side effects to some women. Migration, especially uterine perforations, expulsion, increased menstrual blood loss, pain, and uterus or pelvic inflammatory and risk of extrauterine pregnancy after IUD insertion are frequently occurred on patients [11, 12]. IUD migrated into bladder is a rare and serious complication [13]. On the other hand, the burst release of the cupric ion (Cu^2+^) of cooper containing IUDs (Cu-IUDs) is an important side effect particularly in the first month. The Cu-IUDs, levonorgestrel-releasing intrauterine system (LNG-RIS) and implants are LARC. Excessive Cu^2+^ may cause toxic effect on the cell and increase bleeding and pain [14–16]. The LNG-RIS can increase irregular bleeding and/or spotting days particularly after 3 months of use [17, 18]. In addition, the shape, size and weight of IUDs also have a great impact on contraceptive effects and side effects. The increase in menstrual volume caused by IUD seems to be related to the size of the device. The greater the size and weight of the IUD, the greater the amount of menstrual blood loss is. ^11,12^ Relationships between the size of the IUD and the size of the uterine cavity are also considered to be a factor in the expulsion of the IUD [12].

Inducing local inflammatory reaction in the endometrium is the major effect of all IUDs. The inflammatory response can be enhanced by Cu-IUDs. Cu+ is released from Cu-IUDs and reached in a concentration in the luminal fluids of the genital tract that is toxic for spermatozoa and embryos [19]. Cu-IUDs are usually associated with menorrhagia. As a local foreign body reaction, Cu-IUDs can cause certain morphological changes in the endometrium and infiltration of monocytes as well as a few plasma cells during the proliferative phase of the cycle [20]. The ovulation dysfunction hemorrhage may be related to morphological and biochemical changes in the IUD use. The LNG-RIS has profound morphological effects upon the endometrium, which may lead to a large number of decidualization of endometrial stromal cells, atrophy of the glandular and surface epithelium and vascular morphology change [21, 22]. Irregular bleeding is still a common reason for the discontinuation of progestin-only contraception [21]. Down-regulation of sex steroid receptors is found in all cellular components with endometrial exposure to LNG-RIS [22]. Paired box 2 (PAX2) is a member of paired box family and is an oncogene involved in the development of endometrial cancer. Recent studies have demonstrated that occurrence of PAX2 loss expression in endometrial hyperplasia increases with malignant progression, and PAX2 gene is required for embryonic uterine development, during endometrial carcinogenesis [23–25]. No study has been found in the effect of IUDs as the contraceptive method on endometrial sex steroid receptors and PAX2 expression in uterus of rhesus macaques.

We have reported that the main contraceptive mechanism effect of the three-dimensional reticular IUD (3-DRIUD) in rats, and observed that the larger the physical space occupied by the intrauterine device, the less the pregnancy [26]. In present study, we designed and manufactured a novel three-dimensional grid intrauterine device (3-DGIUD) with nickel-titanium (nitinol) wire and silicone rubber for rhesus macaques, and investigated the changes in the uterus and endometrium of macaques after the 3-DGIUD placement.

## Materials and Methods

### Materials of the new 3-DGIUD and inserter for rhesus macaques

The frame of the 3-DGIUD was composed of nitinol wire (with a diameter of 0.05 mm) and covered with a layer of silicone rubber. Design the structure of the 3-DGIUD was according to the size and shape of the rhesus macaque’s uterus. The shape of the 3-DGIUD for rhesus macaques was three-dimensional in nature and had a reticular grid shape. Its height (H), upper width (D), lower width (d) and thickness were 0.6-1.2 cm, 0.4-0.6 cm and 0.2-0.4 cm, respectively (Figure 1 A, B and D). The weight of the 3-DGIUD was 0.015-0.020 g. A layer of silicon-boron coupling agent was covered on the surface of the 3-DGIUD, and finally multi-layer coating method was used with silica gel, and the 3-DGIUD coated with silica gel was vulcanized. The 3-DGIUD was placed in the external casing tube of the inserter (Figure 1 C). Figure 1 shows photographs of the 3-DGIUD for rhesus macaques.

**Figure 1.**
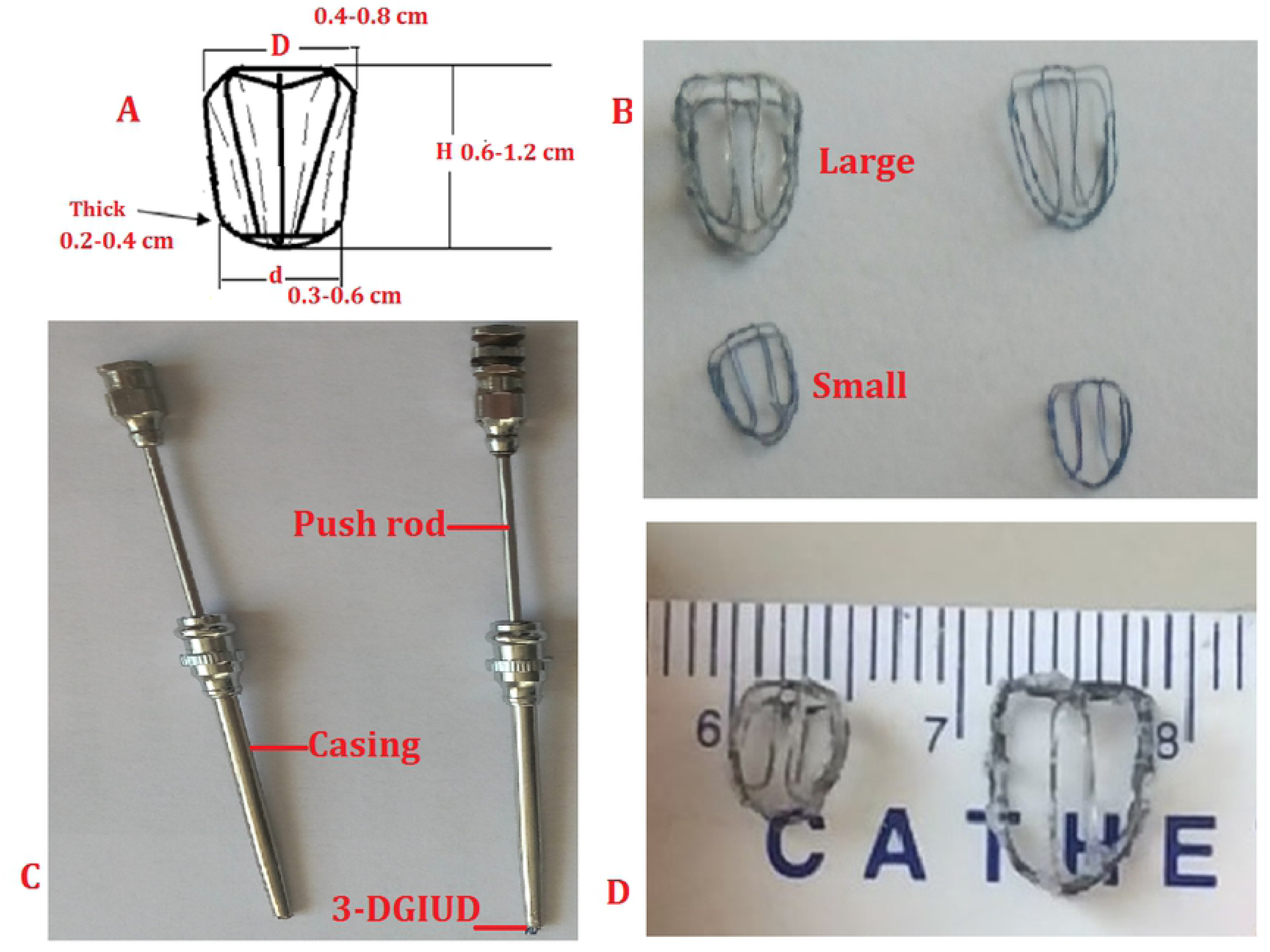
Design and manufacture of the novel three-dimensional grid intrauterine device (3-DGIUD) for rhesus macaques. (A) Designed map of 3-DGIUD. (B) Actual 3-DGIUD, two types, large and small. (C) Inserters with the 3-DGIUD for rhesus macaques. (D) Measured 3-DGIUD size.

The material of the inserter of the 3-DGIUD for rhesus macaques was stainless steel. Inserters of the 3-DGIUD were composed of an external casing tube (diameter 2.0 mm) and an internal needle core (push-rod) (diameter 1.8 mm) (Figure 1 C).

### Animals

Rhesus macaques were from the Fujian Provincial Non-human Primate Animal Experimental Center. All procedures were performed in accordance with “Guidelines for the Care and Use of Experimental Animals in Fujian Province”, and approved by the Animal Care and Use Committee of Fujian Provincial Institute for Family Planning Science and Technology. In December 2016, twenty adult female rhesus macaques were divided into 3 groups, 9 macaques were in the 3-DGIUD group, 9 cases were taken as the sham operation group, and the other 2 cases were taken as control group (without surgery). The license number of the Animal Care and Use was SYXK (Min) 2015-0007 and SCXK (Min) 2015-0002. All macaques were single cage (stainless steel) with standard feeding. The indoor temperature of the animal was 22∼25°C, the relative humidity was 60∼70%, artificial lighting was 12h/d, and the air was ventilated.

### Surgical procedures

The surgical procedures were carried out under sterile technique. Macaques were sedated with ketamine, atropine sulfate (0.02 mL/kg intramuscular injection) and ketamine hydrochloride (0.1–0.2 mL/kg intramuscular injection in a 50 mg/mL aqueous solution), the abdominal regions were shaved and the animals were positioned in the supine position. The lower abdominal region was disinfected with 70% ethanol and iodine tincture and covered with sterilized drapes, and then surgical midline lower abdominal incision was performed and the uterus was exposed (Figure 2 A and B). The external dimensions of the uterus were measured with a sterilized caliper (Figure 2 B). Then a catheter was inserted into the uterus (Figure 2 C). The 3-DGIUD was placed into the uterine cavity with a catheter for 9 macaques (3-DGIUD group). The sham surgery was performed for another 9 macaques (only the catheter inserted into the uterine cavity, without the 3-DGIUD placement). Then the incision of the uterus was sutured with absorbable sutures (Figure 2 D), and macaques received prophylactic antibiotics (Cefazolin, 30 mg/kg).

**Figure 2.**
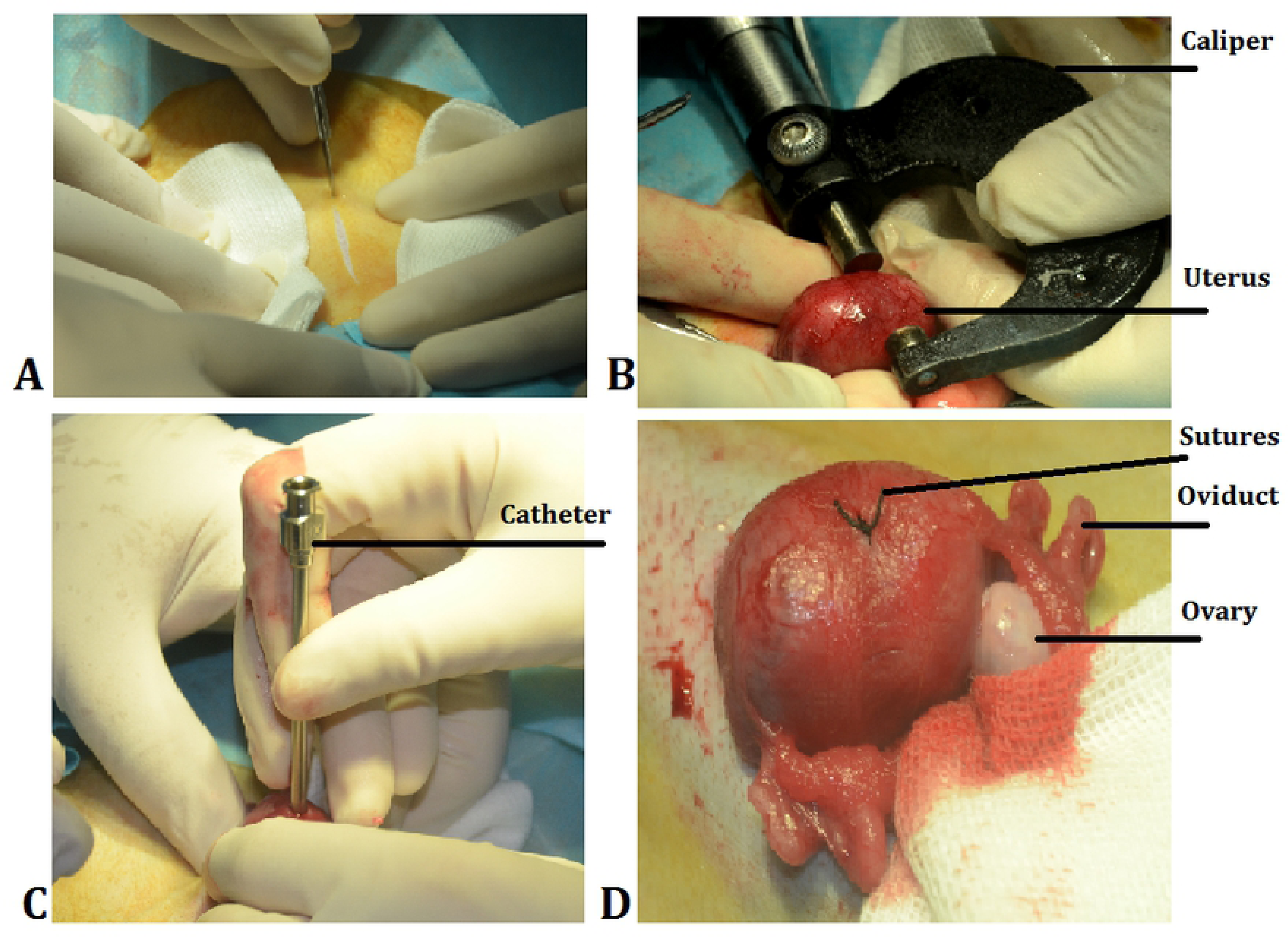
Placement of 3-DGIUD. (a) The lower abdominal incision. (b) Measuring the uterine size. (c) A catheter inserted into the uterine cavity. (d) After placement of 3-DGIUD, the incision in the uterus was sutured.

### Treatment phase and ultrasonography

On the 10^th^-day after operation, 18 female macaques in the 3-DGIUD group (n=9) and the sham operation group (n=9) were coupled with fertile male macaques (1:1) to observe the effect of contraception of the 3-DGIUD. Two macaques in the 3-DGIUD group and one macaque in control group (No. 1, without surgery) were hysterectomized at third-month. On 12 months after the 3-DGIUD placement, 6 macaques in the 3-DGIUD group and 1 macaque in control group (No. 2, without surgery) undergo hysterectomy.

Abdominal ultrasound was performed for 18 macaques in the 3-DGIUD group and the sham operation group every month, to check the uterus, 3-DGIUD and pregnancy. For ultrasonography, we used an X300PE (Siemens, Germany) machine with a 3.5-MHz (VS 13-5, Siemens) probe.

After hysterectomy, the uterus was used for histology and immunohistochemistry and morphological studies. Several cross-sections (about 2 mm thick) were cut freehand from the lumen to the myometrial border with a razor blade under stereomicrocope magnification. These slices were processed further to assess histological development, steroid receptor immunocytochemistry and markers (PAX2) of proliferation. Steroid receptors (estrogen and progestin receptor, ER and PR), and PAX2 described as below.

### Histology and immunohistochemistry

Pathological examination was carried out to the endometrium. Samples for histology were fixed with a mixture of 2% glutaraldehyde and 3% paraformaldehyde, embedded in glycol methacrylate (GMA), sectioned, and stained with hematoxylin and eosin (HE). The expression of ER, PR and PAX2 were detected by immunohistochemistry of endometrium on 12 months after surgery. For immunohistochemistry, briefly, paraffin-embedded tissue sections were de-paraffinized with xylene and dehydrated through graded ethanol, and then their endogenous peroxidase activity was quenched with 3% hydrogen peroxide for 30 min. Antigen retrieval used 10 mM sodium citrate buffer for 2 min. Sections were washed with PBS (phosphate-buffered saline) and blocked with goat serum for 15 min. Sections were incubated with blocking serum for 20 min and then with the primary monoclonal anti-ER (1D-5; detects ER alpha, Biogenex, San Ramon, CA, USA) and anti-PR (JZB-39; courtesy of Geoffrey Greene, University of Chicago, detects PR-A and PR-B), and both were incubated overnight at 4 °C, washed, and then were incubated for 20 min at room temperature with respective biotinylated goat anti-mouse/rabbit secondary antibody and biotinylated horseradish peroxidase complex both in the Ultra Sensitive™ SP (Mouse/Rabbit) IHC Kit (Maixin Bio). For PAX2, Sections were dried for 1 h at 65 °C before treatment procedure of deparaffinization, rehydration and epitope retrieval in the Pre-Treatment Module, PT-LINK (DAKO) at 95 °C for 20 min in 50 × Tris/EDTA buffer, pH 9.0. Before staining the sections, endogenous peroxidase was blocked. The antibodies used were against 6H2.1. After incubation, the reaction was visualized with the stain used: PAX2, clone: Z-RX2. Sections were counter-stained with hematoxylin. Appropriate negative controls including no primary antibody were also tested. The sections were incubated with DAB (3,3’-diaminobenzidine tetrahydrochloride) for 5 min (Maixin Bio), washed under tap water and counterstained with hematoxylin to facilitate identification of cellular elements. The section was cover-slipped. Finally, the slides were observed by microscopy (Olympus).

### Statistical analysis

The statistical analysis of the study was performed using an IBM SPSS (Version 22.0. Armonk, NY: IBM Corp.). Data were shown as the mean ± SD. Parametric data were analyzed statistically using Student’s *t*-tests. The exact Pearson Chi-Square test (Fisher’s Exact Test) was used for the pregnant rate. The difference was considered statistically significant for *P* values <0.05.

## Results

### Uterine corpus measurement results before operation and during surgery

The weight, uterine corpus measurement (during surgery) and ultrasonography (before operation) were shown in Table 1. The mean weight and mean age of the 3-DGIUD group (n=9) and the sham operation group (n=9) were 6.4 ± 1.0 kg and 129.6 ± 1.6 months, and 6.5 ± 0.8 kg and 130.8 ± 8.4 months, respectively, no significant differences were found between both groups. The weight and age of another 2 female control macaques (without operation) were 5.9 kg and 127 months and 6.9 kg and 132 months, respectively. In longitudinal section, the uterus was shaped like an inverted pear (Figure 3 A and B), whereas in transverse section, it was triangular, with a well-defined border by ultrasound. The 3-DGIUD in the uterus shown strong echo, and longitudinal and transverse section were 8 × 6 × 3 mm (large) (Figure 3 C and D) and 6 × 4 × 2 mm (small) (Figure 3 E). The mean size of uterine corpus of height, width and thickness during operation measured by caliper was 28.1 ± 1.0 mm, 21.7 ± 2.5 mm and 17.3 ± 2.6 mm in the 3-DGIUD group, and 28.7 ± 0.8 mm, 22.8 ± 2.8 mm and 17.3 ± 1.6 mm in control group (*t*=0.533, 1.153 and 0.000, and P=0.609, 0.282 and 1.000); the size of uterine corpus by ultrasound in longitudinal section and in transverse section was 28.12 ± 2.26 mm, 19.49 ± 2.11 mm and 21.77 ± 2.43 mm in the 3-DGIUD group, and 29.63 ± 2.25 mm, 19.24 ± 1.67 mm and 21.33 ± 2.27 mm in control group (*t*=1.240, 0.297 and 0.589, and P=0.250, 0.787 and 0.572), respectively.

**Figure 3.**
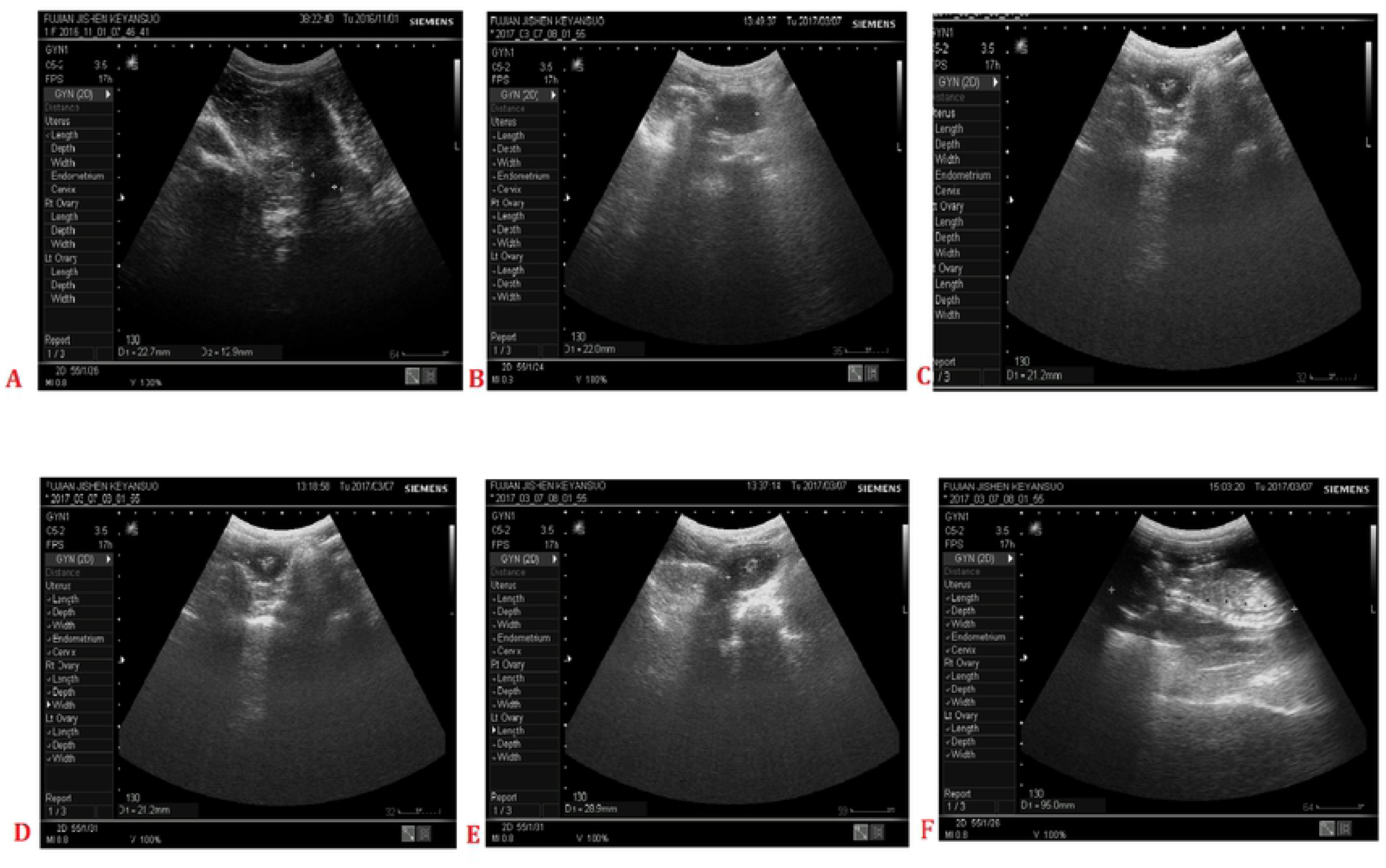
The uterine ultrasound of rhesus macaques. (A) and (B), before surgery, the longitudinal section and the transverse section. (C) and (D), one month and 3 months after surgery, strong echo of the large type of 3-DGIUD. (E), the small type of 3-DGIUD in the uterine cavity. (F), pregnancy, fetal in the uterus.

**Table 1.**
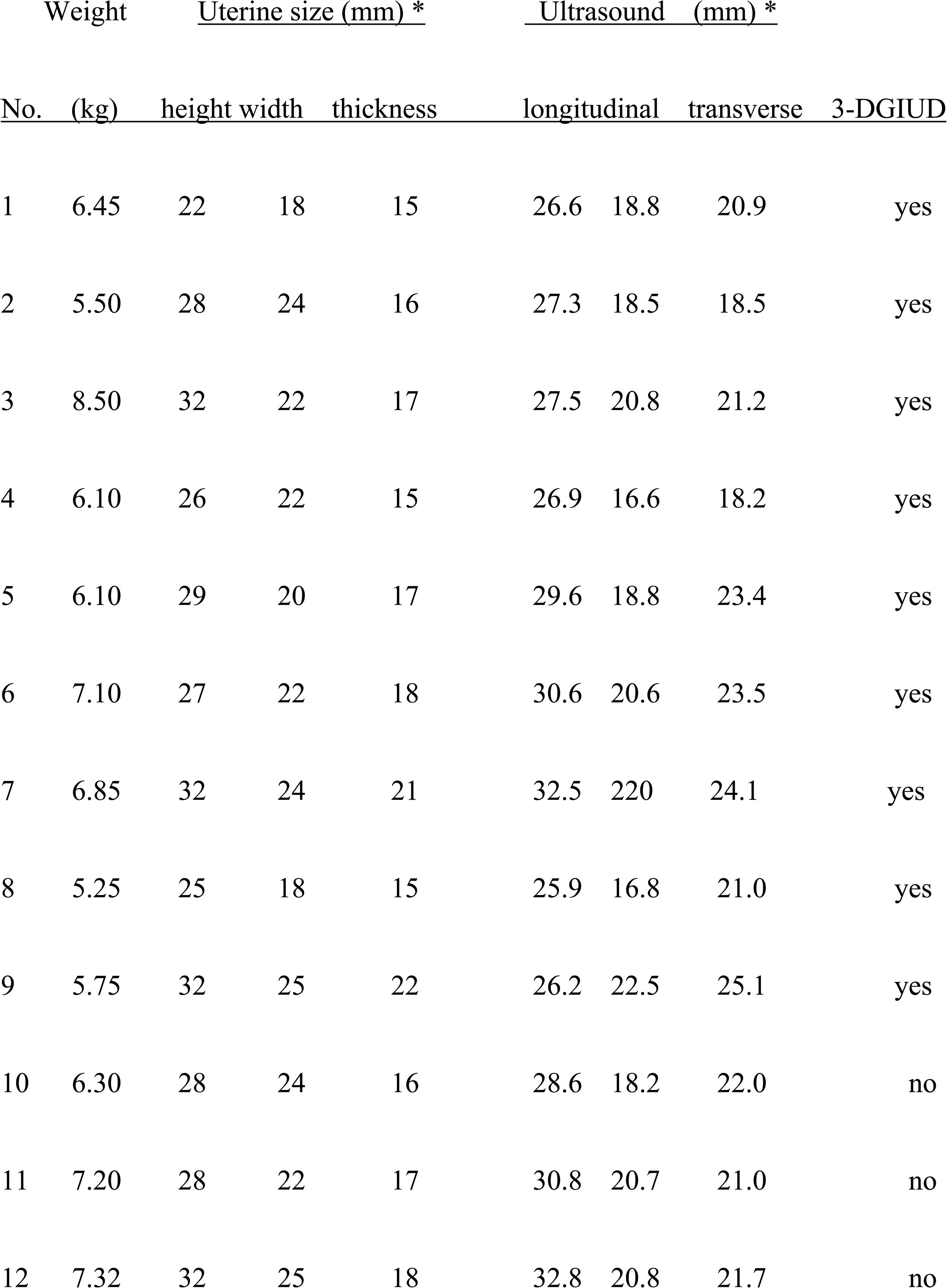
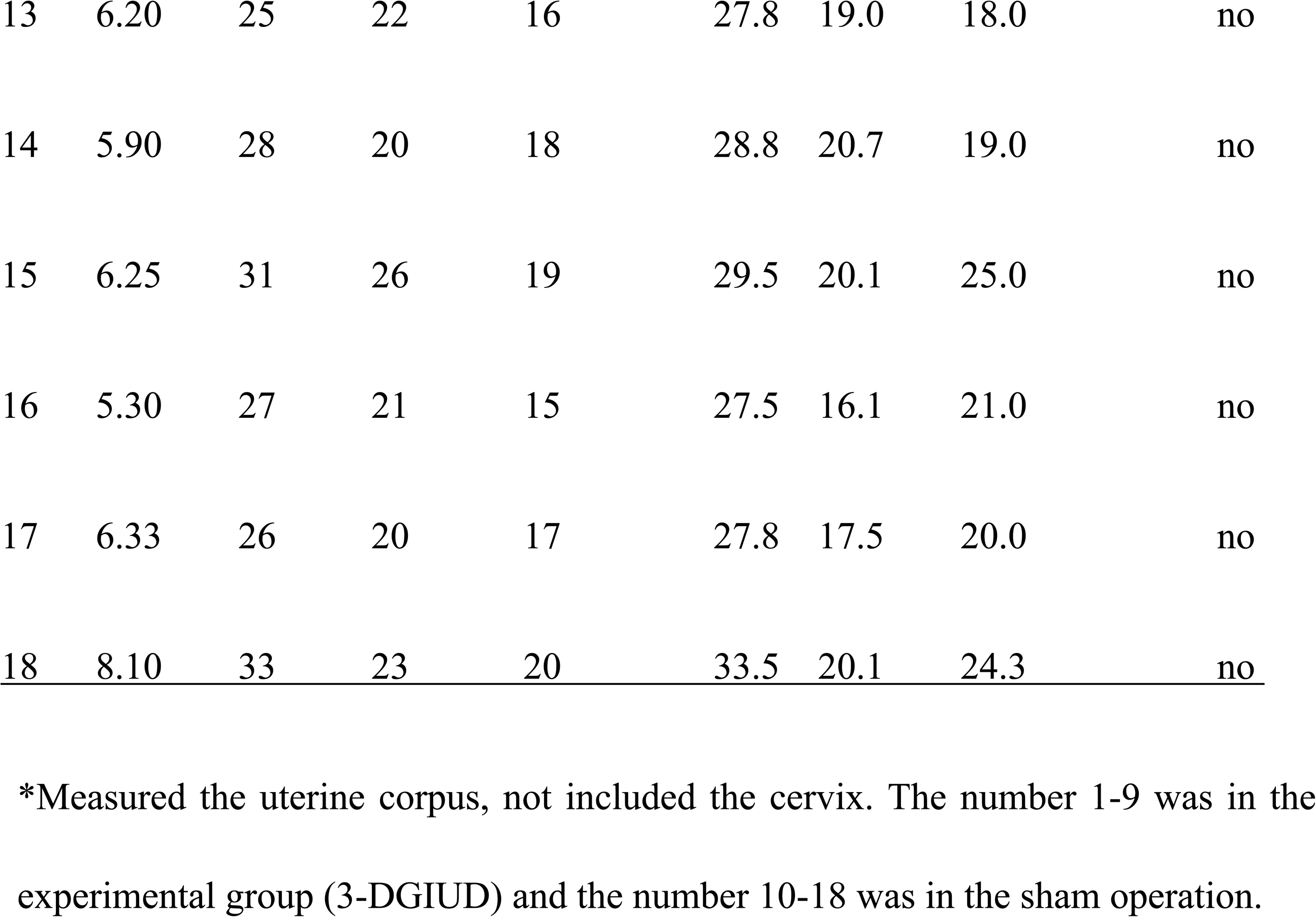
The weight and uterine size by ultrasound before and during surgery

### Pregnancy and uterine measurement of rhesus macaques on 3 months and 12 months after surgery

The large type of the 3-DGIUD was placed in 7 rhesus macaques and the small type was placed in 2 cases. After 3 months, the 3-DGIUD loss and pregnancy were found in one macaque (1/9, 11.1%), and this 3-DGIUD was small type. In the sham operation group, 6 and 3 macaques on 3 and 12 months after surgery were pregnant (Figure 3 F), respectively, and there was significant difference comparing with the 3-DGIUD group (*x*^2^ = 5.844, P = 0.05 and *x*^2^ = 14.400, P = 0.000). Uterine measurement on 3 months and 12 months after surgery is shown in Table 2.

**Table 2.**
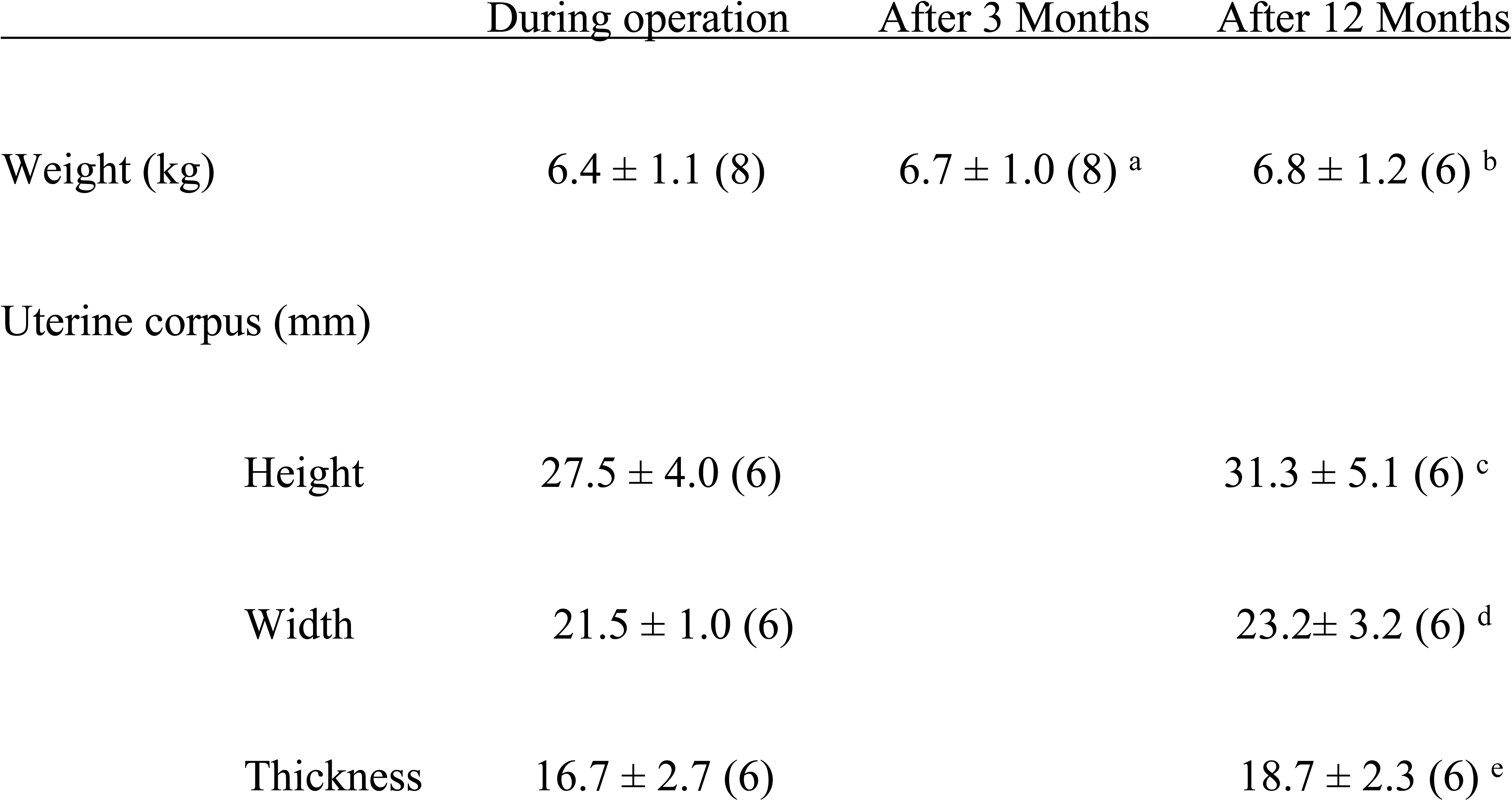
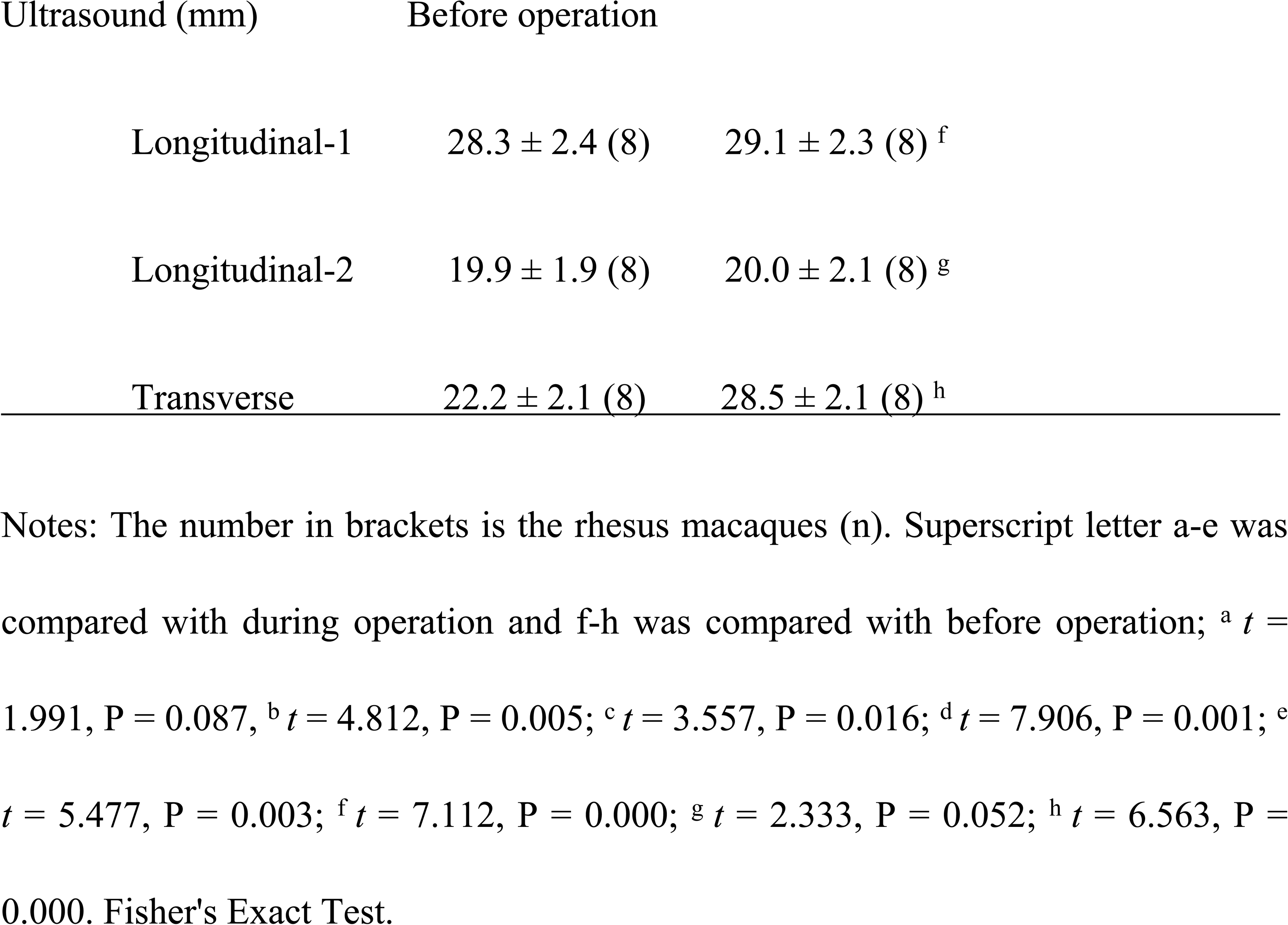
Weight and uterine size of macaques on 3 and 12 months after surgery

### Changes in the histology

No pelvic infection occurred in 18 rhesus macaques on 3 and 12 months after surgery. After hysterectomy, the uterus was cut open. It can be seen that the 3-DGIUD did not adhere to surrounding tissues or embed into myometrium (Figure 4 A). The endometria in the without operation macaque (No. 1, control) were typically straight tubular glands in a normal stroma (Figure 4 B). Pathological examination of uterus on 3 months after the 3-DGIUD placement: Endometrial epithelium was intact (Figure 4 C and D), interstitial cells were edema, short fusiform and dense. A small number of glands were curved and vacuole. The spiral artery was hyperplasia on 3 months after 3-DGIUD placement (Figure 4 C). After the 3-DGIUD placement for 12 months, endometrial monolayer columnar epithelium was mostly intact, focal epithelial cells were loss with focal hemorrhage, and a few neutrophils infiltration were observed. Some glands secreted vacuoles. The vitreous degeneration was found in the basal layer and the superficial muscle layer; and the interstitial spiral arterioles developed well (Figure 4 D).

**Figure 4.**
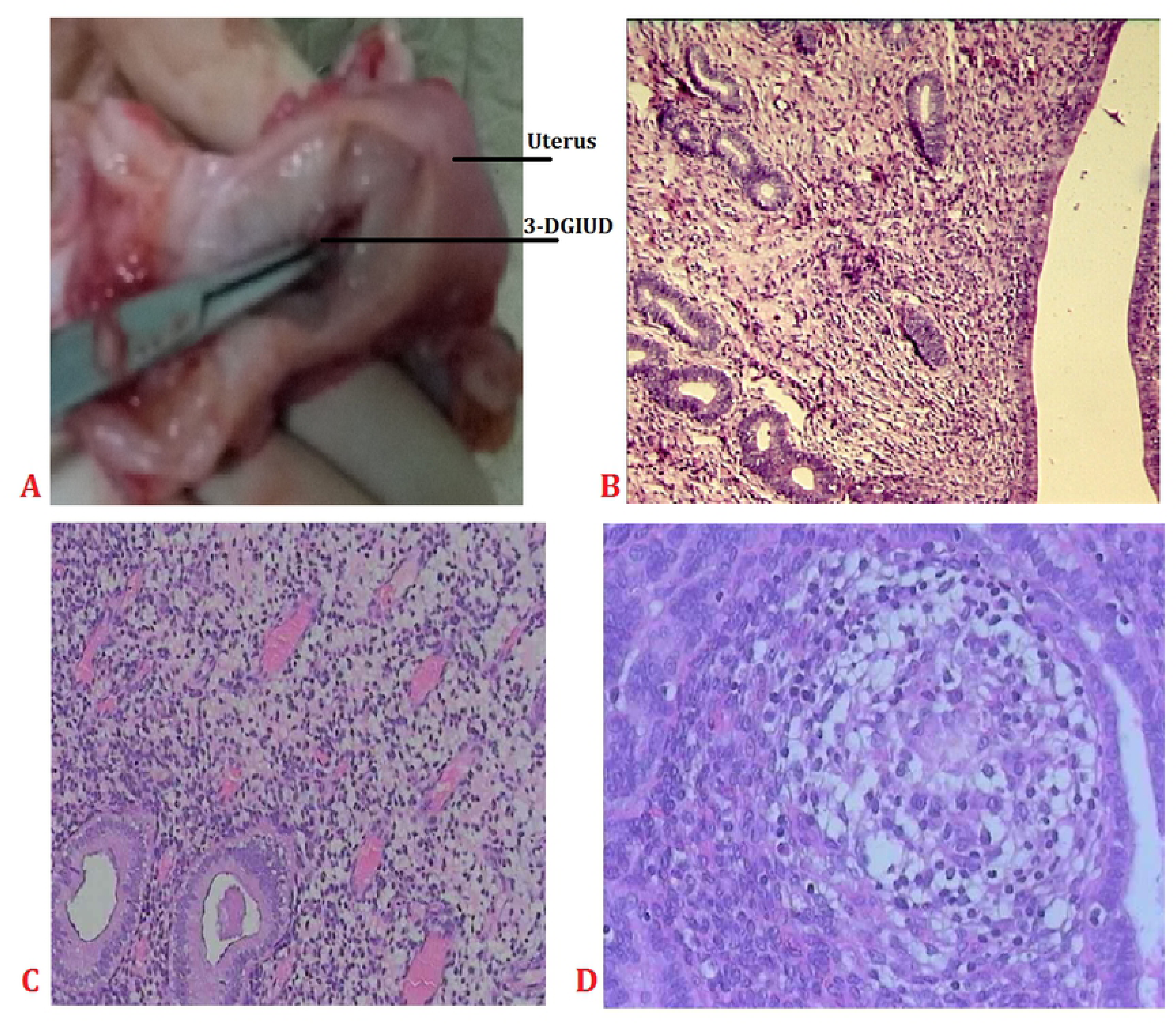
The 3-DGIUD and endometrial histology of macaques. (A), the 3-DGIUD in the uterine cavity. (B), the uterine histology of the control (No.1, non-operation macaque) (20 **×** magnifications). (C), the uterine histology of macaque on 3 months after the 3-DGIUD placement (20 **×** magnifications). (D), the uterine histology of macaque on 12 months after the 3-DGIUD placement (25 **×** magnifications).

### Changes in the immunohistochemistry

There was no difference in cell distribution and staining intensity of the expression of ER, PR and PAX2 between non-surgical rhesus macaque (No.2, control group) and the 3-DGIUD placement group. The Figure 5 illustrates representative examples of the ER, PR and PAX2 immunostaining of uterine sections of normal endometrium (No. 2, control) (Figure 5 A, C and E) and the 3-DGIUD group (Figure 5 B, D and F). The positive ER, PR and PAX2 were shown in the nuclear and cytoplasmic immunostaining of endometrial glands. ER, PR and PAX2 expression status in endometrium was normal (Figure 5 B, D and F). No loss, increase and decrease of ER, PR and PAX2 protein expression were found.

**Figure 5.**
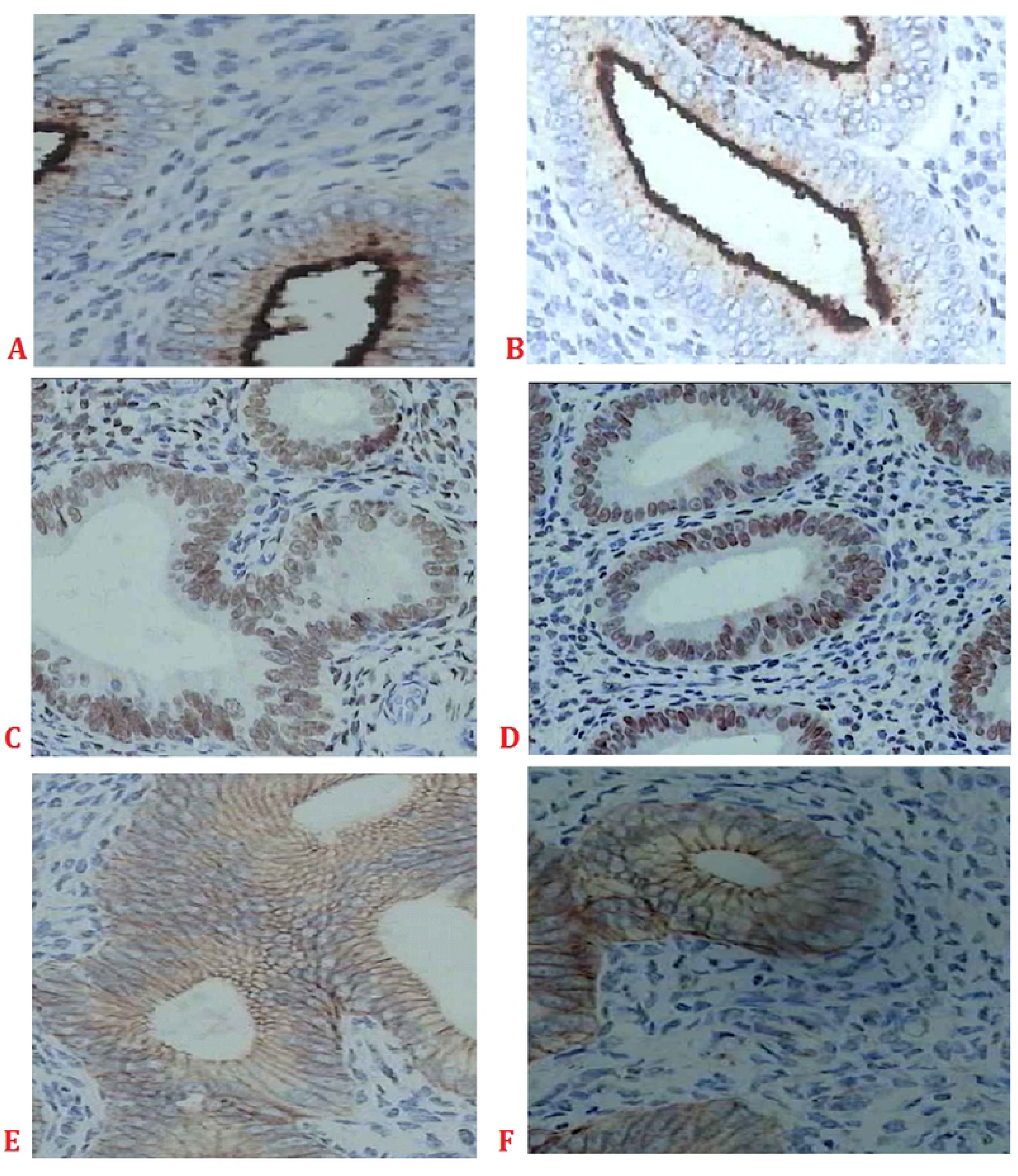
Uterine immunohistochemistry of the ER, PR and PAX2 on 12 months after surgery. (A), (C) and (E), non-operation macaque (No. 2, control). (B), (D) and (F), the 3-DGIUD group macaque. (A) and (B), the ER immunostaining, both were similar. (C) and (D), the PR immunostaining, both were similar. (E) and (F), the PAX2 immunostaining, both were similar. All were 25 **×** magnifications.

## Discussion

### Principal findings of the study

We designed the shape of the 3-DGIUD by using nitinol wire, performed rhesus macaque experiments, and investigated the changes in the uterus of rhesus macaques in this study. The uterine shape and size of the rhesus macaque were similar to human (only smaller than human). Macaques (non-human primate animal) may be the best animal model for experiment of IUDs. We explored that the weight and uterine size of macaques increased on 3 months and 12 months after the 3-DGIUD placement. Changes in endometrial epithelium by the local oppression of the 3-DGIUD were observed. No alterations were found in the expression of endometrial ER, PR and PAX2 after the 3-DGIUD placement.

### Results of the study in the context of other observations

#### Materials of the frame of IUDs

The frame of currently used IUD is made of stainless steel and plastic, and included copper or additional hormones [27, 28]. The Cu (CuT380A) IUD is the only non-hormonal LARC device approved by the United States Food and Drug Administration (FDA) [29]. The Cu^2+^ of Cu-IUD-releasing such as small T-shaped devices which made of flexible plastic, can kill sperm. The Cu-IUD is a LARC (up to twelve years), however some women discontinue use due to undesired side effects such as pain or cramping and complaints of heavy bleeding [14-16,28]. The high rate of amenorrhea is often seen with the higher progestin devices [30, 31]. In present study, the frame of the new 3-DGIUD is made of nitinol wire and silicone rubber. The nitinol wire is no toxic and used widely in clinical such as orthopedics, bone and cardiovascular stents. The nitinol frame was covered with silicone rubber, to prevent the 3-DGIUD from adhering to the endometrium. The metal copper and progestin are not used for the 3-DGIUD. These may avoid side effect of copper and hormone IUDs.

#### Efficacy and side effects of IUDs

Proper installation of IUD or IUD system will reduce adverse effects and improve acceptability, resulting in enhanced continuation of the IUD use [32]. Cramping pain, erratic bleeding or menorrhagia and expulsion of IUDs may be caused by the dimensional incompatibility. In the present study, the small type of 3-DGIUD was expulsion in one macaque at 3 months. The remaining 8 macaques, no pregnancy occurred after surgery. Therefore, the size of IUD is too small, easy to fall off, leading to pregnancy. On the other hand, menorrhagia, dislocation or expulsion and contraceptive efficacy may be affected by the shape and the weight of IUDs. If the shape is too large or the weight is too heavy of the IUD, severe compression will be caused. The rigidity of the inserted tube may also be linked to risks [33, 34]. In the present study, we improved the shape of IUDs. The three-dimensional structure for macaques replaced the two-dimensional structure of the commonly used IUDs. The weight of the 3-DGIUD’s frame was light. The space of uterine cavity was occupied by the 3-DGIUD and embryo implantation was interfered. Nitinol has a memory function, and can restore to the designed shape at body temperature. It has a good flexibility, may be conformed to the contraction and activity of the uterus, and avoided uterine perforation.

The contraceptive efficacy is very important for IUDs. Wu *et al.* reported [35] that the LNG-IUS placed in 3 monkeys, expulsion of device is found in one monkey. In human women, the expulsion rate of postpartum IUD varies according to the placement time, delivery method, and the type of IUDs, ranging from 1.9% to 29.7% [36–39], while removal rates are 3.6% to 19.3% due to associate side effects (bleeding, pain and discharge) [36, 38].

### Clinical implications of the study

#### Effects of IUDs on the uterus

Wang et al. [40] reported that the chronic endometrial inflammation of histologic features occurs after placement a bare copper wire to contraception in the uterus of rhesus macaques. The chronic and non-specific endometrial inflammation may be one of contraceptive effects of the Cu-IUD. The strong local inflammatory response is induced by LNG-IUS for the transplant recipients as in the healthy control [41].

#### The detection of ER, PR and PAX2 expression can be used to predict the response of endometrial hyperplasia and cancer for IUD use

Two studies reported that IUD has nothing to do with the increased risk of breast cancer [42, 43], whereas other studies reported that IUDs are associated with an increased risk of breast cancer [44, 45]. In the LNG-IUS used patients, the recurrence and formation of endometrial polyp may be inhibited through lowering the expression of ER and PR [46]. The LNG-IUS can reduce the expression of ER and PR in endometrium and inhibit endometrial proliferation [47]. When the conservative treatment with LNG-IUS failed, the expression of PR and ER of these patients were higher [48]. The complete down-regulation of PR and ER expression in uterine glands and stroma is caused by the LNG-IUS in human endometrial hyperplasia [49]. The LNG released locally from the IUD has a depressive effect on the ER and PR, which may contribute to the contraceptive effect of this type of IUD and may also be the causes of LNG-IUS-induced irregular bleeding and amenorrhoea [50]. In this study, PAX2 expression was normal, and there was no decrease, loss or over-expression after placement 3-DGIUD. PAX2 is a downstream gene in the steroid hormone receptor signal pathway. It is over-expressed in endometrial cancer and benign endometrial hyperplasia [51]. Recently, Monte et al. [23] and Quick et al. [25] reported that PAX2 deficiency in up to 77% of endometrial adenocarcinoma and 71% of patients with atypical endometrial hyperplasia. As an oncogene involves in the development of endometrial cancer, the expression of PAX2 is increased in the neoplastic lesion progresses from a premalignant state to endometrial cancer. Knock-down of PAX2 may lead to the decrease of cell viability, invasion and migration, while PAX2 over-expression causes to the opposite effects. PAX2 acts as a tumor suppressor in proliferative and self-renewing endometrial epithelial cells [23].

### Research implications

#### Unanswered questions

Birth control plays pivotal roles in the reduction of maternal, infant, and child mortality. As the main method of contraception, IUD is the focus of clinical research. Questions relating to the risks of IUD use remain unanswered. The material, size and shape of the IUD have significant impacts on contraceptive efficacy, and side effects may be avoided or decreased by changes in the shape and materials of IUDs. The size of IUDs is too small to be expelled [33, 34]. In the present study, a small type of 3-DGIUD was expelled from the uterus after 3 months.

#### Proposals for future research

Contraceptive effectiveness and side effects of IUDs should be studied by improvement of the material and the shape of IUDs. The material IUDs must be non-toxic and to fit for the uterus.

## Conclusion

In conclusion, despite the 3-DGIUD was developed for macaques, it is likely that improvement of the shape and the materials for currently used IUD of human according to this study. The 3-DGIUD was non-toxic and had good contraceptive effectiveness for macaques.

## Data Availability

All relevant data are within the paper and its Supporting Information files.

